# Microbes are potential key players in the evolution of life histories and aging in *Caenorhabditis elegans*

**DOI:** 10.1101/2022.02.26.482130

**Authors:** Josiane Santos, Margarida Matos, Thomas Flatt, Ivo M Chelo

## Abstract

Microbes can have profound effects on host fitness and health and the appearance of late-onset diseases. Host-microbe interactions thus represent a major environmental context for healthy aging of the host and might also mediate trade-offs between life-history traits in the evolution of host senescence. Here, we have used the nematode *Caenorhabditis elegans* to study how host-microbe interactions may modulate the evolution of life histories and aging. We first characterized the effects of two non-pathogenic and one pathogenic *Escherichia coli* strains, together with the pathogenic *Serratia marcescens* DB11 strain, on population growth rates and survival of *C. elegans* from five different genetic backgrounds. We then focused on an outbred *C. elegans* population, to understand if microbe-specific effects on the reproductive schedule and in traits such as developmental rate and survival were also expressed in the presence of males and standing genetic variation, which could be relevant for the evolution of *C. elegans* and other nematode species in nature. Our results show that host-microbe interactions have a substantial host-genotype-dependent impact on the reproductive aging and survival of the nematode host. Although both pathogenic bacteria reduced host survival in comparison with benign strains, they differed in how they affected other host traits. Host fertility and population growth rate were affected by *S. marcescens* DB11 only during early adulthood, whereas this occurred at later ages with the pathogenic *E. coli* IAI1. In both cases, these effects were largely dependent on the host genotypes. Given such microbe-specific genotypic differences in host life history, we predict that the evolution of reproductive schedules and senescence might be critically contingent on host-microbe interactions in nature.

## Introduction

Microbes are thought to have major effects on the evolution and speciation of host populations due to their ubiquitous presence and ability to influence host physiology and health (Bordenstein et al. 2001; Zilber-Rosenberg and Rosenberg 2008; McFall-Ngai et al. 2013). While microbes are best known for their pathogenic or mutualistic effects, they can also modulate how hosts perceive and respond to stressful conditions. This has been observed, for example, in contexts as diverse as viral infections (Martinez et al. 2014) and other biotic stresses (Zhang et al. 2021), the autoimmune response (Langan et al. 2019), drug therapy (Pryor et al. 2019), metabolic dysfunction (Ussar et al. 2016), exposure to high temperatures (Xie et al. 2013; Howells et al. 2016), and chemical toxicity (Coryell et al. 2018). Microbes can thus impact the adaptation of host populations to conditions that are unrelated to the host-microbe interaction itself (Martinez et al. 2016; Faria et al. 2016; Bates et al. 2021; Hoang et al. 2021), which suggests they can also have an indirect and still poorly understood, but fundamental, role in shaping the evolution of host life history and aging.

The progressive loss of physiological function leading to a decline in fecundity and increased mortality, which defines aging, can be explained by the reduced efficacy of selection in purging mutations that have deleterious effects late in life (Fisher 1930; Haldane 1941; Medawar 1946, 1952; Williams 1957; Hamilton 1966; Rose 1991; Kirkwood and Austad 2000; Flatt and Schmidt 2009; Flatt and Partridge 2018). A major mechanism underlying the evolution of aging is antagonistic pleiotropy, i.e., the existence of alleles with antagonistic effects on early and late life-history traits, which lead to genetic trade-offs between fitness components (Medawar 1946, 1952; Williams 1957; Stearns 1989; Rose 1991; Flatt and Promislow 2007; Flatt 2020). Under this model, aging evolves because strong selection for beneficial fitness effects early in life outweighs the deleterious effects of these alleles late in life when selection is weak (e.g., Williams 1957). A large body of work in numerous organisms, including the nematode worm *Caenorhabditis elegans* (Anderson et al. 2011), the fruit fly *Drosophila melanogaster* (reviewed in Flatt 2020), or the fish *Poecilia reticulata* (Reznick et al. 1990), has revealed antagonistic pleiotropy underlying trade-offs by showing correlated responses to selection in major fitness components such as developmental rate, early- and late-fecundity, and lifespan.

Even when populations harbor genetic variation at antagonistically pleiotropic loci, environmental factors may prevent the expression of phenotypic trade-offs and correlated changes in life-history traits (Giesel et al. 1982; Stearns 1989; Ackermann et al. 2001; Sgró and Hoffmann 2004; Gutteling et al. 2007; Swanson et al. 2016). Microbes are likely to be important environmental components in the evolution of aging, given their known effects on host life-history traits (Little et al. 2002; Decaestecker et al. 2003; Brummel et al. 2004; Vale and Little 2012; Leroy et al. 2012; Laughton et al. 2014; Parker et al. 2014; Diaz et al. 2015; Zurowski et al. 2020) and their evolution (Sorci and Colbert 1995; Gibson et al. 2015; Walters et al. 2020). Causal relationships between the composition of the intestinal microbiome and aging observed in humans (Claesson et al. 2011) and other organisms (Clark et al. 2015; Sonowal et al. 2017; Bárcena et al. 2019) are consistent with this idea.

Studies with the *C. elegans* model hold great promise for an improved understanding of the interplay between host-microbe interactions and the evolution of aging. For example, the worm system has been extensively used in the identification of the genetic pathways underpinning aging and longevity (Garsin et al. 2003; Kurz and Tan 2004; Antebi 2007; Evans et al. 2008; Leroy et al. 2012), many of which are shared with humans (Kurz and Tan 2004). At the same time, *C. elegans* has also been a valuable tool for studying host-microbe interactions (e.g. Tan et al. 1999; Abbalay et al. 2000; Garsin et al. 2003; Schulenburg et al. 2004; Coolon et al. 2009; Leroy et al. 2012; Diaz et al. 2015; Dirksen et al. 2016; Schulenburg and Félix 2017; Zhang et al. 2021) and how, either through the nutritional content of bacteria or specific pathogenic effects, such interactions regulate host development, reproduction, metabolism, immunity, and lifespan (MacNeil et al. 2013; Pang and Curran 2014; Chan et al. 2019). Notably, links between immunity and aging are well established in *C. elegans* (Garsin et al. 2003; Kurz and Tan 2004; Troemel et al. 2006; Evans et al. 2008), for example in the context of lifespan expansion obtained with specific bacterial metabolites (Virk et al. 2012; Gusarov et al. 2013; Han et al. 2017) or by transferring worms from their regular food source (*Escherichia coli* OP50) to other bacteria such as *Bacillus subtilis* (Aballay *et al*. 2000; Portal-Celhay et al. 2012; Donato et al. 2017).

In support of the importance of host-microbe interactions in the evolution of *C. elegans* in its natural settings (Félix and Braendle 2010; Schulenburg and Ewbank 2014; Martin et al. 2017), microbial effects have been shown to vary between *C. elegans* genotypes, represented by different wild type strains (Schulenburg and Ewbank 2004; Martin et al. 2017; Zhang et al. 2021). Interestingly, the worm’s genotype was also shown to have an active role in determining the gut colonization success of different bacteria (Dirksen et al. 2016; Zhang et al. 2021).

To date, it remains largely unclear to what extent the evolution of life histories and senescence in nematode hosts depends on specific host-microbe interactions. To address this question, we studied the impact of different pathogenic and non-pathogenic bacteria on the reproductive schedule and survival of *C. elegans*. To this end, we focused on two non-pathogenic *E. coli* strains, a pathogenic *E. coli* strain, and a pathogenic *Serratia marcescens* strain. First, we confirmed that the effects of each microbe on the host’s reproductive timing and lifespan depended on the host’s genotype, suggesting the potential for local adaptation to the microbial environment. Secondly, we studied a genetically diverse, male-female (gonochoristic) laboratory-derived *C. elegans* population (Theologidis et al. 2014) to study how microbes affect life-history traits (age-specific and total fertility, age at first reproduction, male and female developmental rate, lifespan) and their evolution in *C. elegans*.

## Material and Methods

### BACTERIAL STRAINS

Bacterial strains used in our experiments included two commonly employed non-pathogenic *Escherichia coli* strains, OP50 (Brenner 1974) and HT115(DE3) (Timmons et al. 2001), and two pathogenic strains, E. coli IAI1 (Picard et al. 1999; Diard et al. 2007) and *Serratia marcescens* Db11 (Flyg et al. 1980; Kurz et al. 2003)*. E. coli* HT115(DE3) had been used as food during the establishment of the *C. elegans* D00 population described below. The strains *E. coli* HT115(DE3), *E. coli* OP50, and *S. marcescens* Db11 were obtained from the Caenorhabditis Genetics Center (CGC), and the *E. coli* IAI1 strain was kindly provided by Ivan Matic.

### NEMATODE POPULATIONS

To assay life-history responses to the four above-mentioned microbe strains we used the N2 lab-adapted strain and 4 wild isolates (CB4852, CB4855, CB4856, PX174), each one being isogenic with respect to a different genotype (from here on designated ‘individual genotypes’), and the outbred experimental *C. elegans* population, D00. The D00 population was first described by Theologidis et al. (2014), being a genetically diverse dioecious population (with males and females) established by introgression of the *fog-2(q71)* mutant allele (Schedl and Kimble 1988) into the genetic background of a previously laboratory-adapted androdioecious population (consisting of males and hermaphrodites; Teotónio et al. 2012; Chelo and Teotónio 2013). Throughout laboratory adaptation, D00 worms were provided with *E. coli* HT115(DE3) as a food source and the population evolved under discrete (non-overlapping) generations imposed by a 4-day life-cycle, herein referred to as “early reproduction”. This population, characterized by obligate outcrossing, harbors a large amount of genetic variation as a result of an initial mixture of 16 isogenic strains, chosen to represent a significant proportion of the known genetic diversity in *C. elegans* (Rockman and Kruglyak 2009; Teotónio et al. 2012; Noble et al. 2019). The 5 wild strains analyzed in this work were part of that initial mixture.

### GROWTH CONDITIONS

Bacteria were grown overnight in NGM-lite solid media at 37 °C from LB-grown cultures. Nematode maintenance followed previously described protocols (Stiernagle 1999; Chelo 2014). On day one, L1 larvae were seeded on NGM-lite supplemented with ampicillin (100 mg/ml), carrying a confluent lawn of *E. coli* HT115(DE3). 10^3^ larvae were used per plate, and development proceeded at 20°C and 80% (RH) for 72 hours, until day four of the life-cycle. Plates were washed with M9 buffer and a KOH:NaCIO solution was added (“bleaching”) to kill adults and larvae but allowing unhatched embryos to survive. Eclosion of first-stage larvae (L1) occurred overnight in 4 ml of M9 buffer with 2.5 mg/ml of tetracycline under constant shaking.

### POPULATION GROWTH RATES

To understand how reproductive timing is affected by the different bacteria, population growth rates were measured at two different times: at 72 hours after L1 seed (transition from day 3 to day 4), i.e., within hours of reaching sexual maturity (“early reproduction”; Anderson et al. 2011) and at 114 hours post-seed (day 5; referred to as “delayed reproduction”). Frozen populations were thawed and maintained for two generations under standard maintenance conditions, plus one generation in presence of each bacterial strain for acclimatization. In the fourth generation, L1 larvae were seeded on NGM-lite plates (10³/plate) with a lawn of each bacterial strain and allowed to develop for 72 or 114 hours. Following our standard maintenance protocol, cultures were bleached and the number of the live L1s was estimated the following day. Each estimate was obtained by pooling individuals from three plates. Each of the five strains (N2, CB4852, CB4855, CB4856, PX174) and the D00 population were assayed in independent experimental blocks. In the assays, each block included the N2 strain feeding on *E. coli* HT115(DE3) as a common reference, the four different bacteria and the two time points. For each bacterial strain and each time point, we used five technical replicates for D00 and N2 and four technical replicates for each of the other four strains. Data can be found in Supplementary Tables 1 and 2.

### SURVIVAL OF INDIVIDUAL GENOTYPES

The effect of the four bacterial strains on survival was assayed for each of the five *C. elegans* strains (N2, CB4852, CB4855, CB4856, PX174). After thaw and growth for two generations under standard maintenance conditions, L1 larvae were seeded on NGM-lite media (10^3^ individuals/plate) with a lawn of each of the four bacteria. 48 hours later (day 3), L4 hermaphrodites were placed on 24-well NGM-lite plates (five individuals per well), with the corresponding bacteria, which had been grown from a 5 μl inoculum. Individuals were transferred to fresh medium every 24 hours until all were found dead or considered to be missing. Monitoring of missing or dead females occurred at the time of transfer, and individuals were considered dead in the absence of movement or response when being gently touched with a platinum wire. Each of the four non-N2 *C. elegans* strains was assayed in a different experimental block, which also included N2 as a common reference. Four plates were used per block, and every plate included all four bacterial strains. Both the N2 and one of the non-N2 strains were used in every plate, with N2 individuals occupying one fourth of the total number of wells. This experimental design enabled the estimation of plate effects within a block. In total, 480 individuals were assayed in each block, with 120 being N2 individuals and 360 individuals from one of the other isogenic strains. Data can be found in Supplementary Table 3.

### REPRODUCTIVE SCHEDULE AND SURVIVAL OF THE D00 POPULATION

Daily offspring number and survival were monitored to study the effects of different bacteria on individuals of the D00 population. Frozen (−80 °C) stock populations were thawed and maintained for two generations prior to the assay. To set up the experiment, 10³ L1 individuals were seeded on NGM-lite plates carrying each of the four bacteria and incubated until the beginning of day 3 (48 hours later). From each plate, 30 female larvae were placed (one larva per well) onto 24-well plates with antibiotic-free NGM-lite and matching bacteria, as described for the individual genotypes (see above). Adult males from the same population and conditions, but which had been developing for one extra day, were added to the wells (two males per well). Individuals were transferred to fresh medium every 12 hours until day 6, and every 48 hours after day 6, until all individuals were found dead or considered to be missing. During the first five days, males that had died (or were missing) were replaced to ensure mating and fertilization. After removal of adults, plates were kept in the incubator for one day and then transferred to 4 °C for a maximum of two days before counting L2-L3 larvae under the stereoscope with 10x-30x magnification. These data were used to determine total fertility (lifetime reproductive success, LRS), variation in fertility through time and the age at first reproduction (AFR). Survival was scored based on daily observations during the entire period of the experiment, with similar monitoring of missing or dead females as with the individual genotypes. Data can be found in Supplementary Table 4.

### DEVELOPMENTAL RATE OF THE D00 POPULATION

The percentage of individuals that had reached adulthood at a specific chronological time was used as a measure of the developmental rate of the D00 population, with each bacterium. Initial population manipulation followed the protocol for estimation of population growth rates, with two generations feeding on *E. coli* HT115, followed by one generation on each specific bacterium. In the fourth generation, 48h after L1 seeding (1 day prior to the “early reproduction” time), individuals were removed from plates with M9 buffer, washed one time with M9 buffer, centrifuged, and 2 μl from the pellet were placed between a microscope slide and slide cover. Image acquisition was done with a DFK 23UX174 color camera (The ImagingSource) at 10 pixel/μm (60x magnification) mounted in a Nikon SMZ18 stereoscope. ImageJ was then used for manual image analysis to identify the sex of individuals (males and females) and their developmental stages, as L3, L4 or adults, by recognizing morphological distinctive features (state of vulval development and presence of embryos inside the adult for females and tail development for males). Measurements were taken from three experimental blocks with all the four different bacteria being used in each block. This resulted in a mean of 79 +/-21 (SD) individuals being used per bacteria and block combination. Data can be found in Supplementary Table 5.

### DATA ANALYSIS

Statistical analyses were performed in *R* (*R* Core Team 2019). Supplementary files with analyses and *R* code can be found at *FigShare* (see 10.6084/m9.figshare.15022566 for Supplementary Figures and Tables; and 10.6084/m9.figshare.15022599 for Supplementary Data and analysis scripts).

Analysis of population growth rate was carried out using the natural logarithm (ln) of the observed rates. Whenever L1 larvae could not be detected, which would lead to growth rate estimates of zero (two samples; see Supplementary Table 1), values were replaced assuming that one L1 had been observed. To standardize the different blocks with *C. elegans* strains, the growth rates of *C. elegans* N2 with *E. coli* HT115(DE3) were first estimated in each block and at each time point with a random-effects model using a block-specific baseline. The following model was implemented in *R*: *log(GrowthRate) ∼ Time * Bacteria * Celegans, offset=Block_offset*, where *GrowthRate* is the observed L1 growth rate in consecutive generations, *Time* is the number of hours since L1 seed, *Bacteria* represents the bacterial strains, *Celegans* represents the 5 different strains, and *Block_offset* is the value of the block effects obtained with N2 and *E. coli* HT115(DE3).

Cox regression (proportional hazards analysis; Cox 1972) was used to test for differences in survivorship, with *E. coli* HT115(DE3) defining the baseline risk, and assuming right-censored data. For the survival of *C. elegans* strains, mixed-effect models were used with the *coxme* function in *R* (Thernau 2020) in order to include plate effects. The following model was used: *Surv (S.time,S.event) ∼ Bacteria * Celegans + (1|Plate) + Block_offset* (see above). Mean lifespan values based on Kaplan-Meier estimation (Kaplan and Meier 1958) were corrected by the values obtained for each block with N2 (see Supplementary Fig. 1). For the analysis of the D00 population data, the following model was implemented with the functions *Surv* and *coxph* in the *survival* package in *R* (Therneau 2015): *Surv(S.time,S.event) ∼ Bacteria,* with *S.time* being the time at which an individual was found dead or missing (*S.event*). Kaplan-Meier estimation was used to obtain survival curves (see Supplementary Fig. 2) and mean lifespan. Phenotypic association between the growth rate and mean lifespan of individual genotypes were tested with *cor.test* function, independently with each bacterial strain.

For fertility data of the D00 population, observations of 12h intervals were collapsed into daily measures until day 6 and into a single bin beyond that time. Thus, fertility reported for day 3 refers to embryos laid between 48 h and 72 h post-L1 seed, between 72 h to 96 h for day 4, between 96 h to 120 h for day 5, between 120 h and 144 h for day 6, and 144 h onwards to “day 7”. Model fitting and model comparisons were performed with generalized linear models with appropriate error distributions (see below), and analysis of deviance was used to test for significance. Parameter estimates were retrieved and tested with *emmeans* and *pairs* function (Lenth 2018). For pairwise comparisons, we used Tukey’s post-hoc tests and report adjusted *p*-values. The reproductive schedule of the D00 population was modeled with a negative binomial distribution using the *R* function *glm.nb* in the MASS package. The following model was used: *Fertility ∼ Bacteria * Time*, where *Fertility* refers to the number of larvae observed per individual worm during a 24 h period, *Bacteria* represents the four bacterial strains tested, and *Time* is a categorical variable with 5 levels representing the day since the experimental set-up. Post-hoc comparisons were performed between fertility means within each day. Total fertility was modeled with a Poisson distribution using the *glm* function, as follows: *LRS ∼ Bacteria, family = “poisson”(link=”log”),* where *LRS* is the total number of observed larvae. A Gaussian fit was used to analyze AFR with the following code: *AFR ∼ Bacteria, family =”gaussian”,* where AFR (age at first reproduction) refers to the time between L1 seed and the time at which offspring was first observed.

Analysis of developmental rate was done by estimating the proportion of individuals that had reached adulthood at the time measurements took place. For this purpose, individuals identified as L3 or L4 larvae were merged into a single class of “non-adults”. The following generalized linear model was then implemented with the *glm* function in *R*: *Adult ∼ Bacteria * Sex, family =”binomial”*, where *Adult* includes the numbers of adults and non-adults observed with each bacterium, *Bacteria* represents the four bacterial strains tested and *Sex* is either male or female. Analysis of deviance was used to test for significance and Tukey’s post-hoc tests were used for pairwise comparisons.

## Results

### REPRODUCTIVE TIMING AND LIFESPAN OF INDIVIDUAL GENOTYPES

Measurements of population growth rates of the five different *C. elegans* genotypes (Fig. 1A) show an overall decline with time (likelihood ratio test, LRT = 369.4, df = 1, *p <* 0.0001), which depends on the bacteria present in the environment (LRT = 67.57, df = 3, *p <* 0.0001, on the interaction term). In the extreme case, with *E. coli* IAI1, the growth rate decreases an average of 0.41 (± 0.03 SE) per hour. This result contrasts with observations done with *S. marcescens* Db11, where a much lower overall decrease is obtained (estimated slope of −0.04 ± 0.03 per hour). The effect of time is also strongly conditioned on the *C. elegans* genotype (LRT = 38.42, df = 4, *p <* 0.0001, on the interaction term), such that there is a prevalent crossing of the different reaction norms (Fig. 1A). Interestingly, in addition to this overall pattern, genotype-by-time effects are unique within each bacterium (significant three-way interaction between *Time x Bacteria* x *C. elegans,* LRT = 17.71, df = 12, *p* < 0.0001, see methods) and can result in unexpected patterns, as with the CB4855 genotype, which shows an increase of population growth rate between 72h and 114h exclusively in presence of *S. marcescens* Db11.

**Figure 1.**
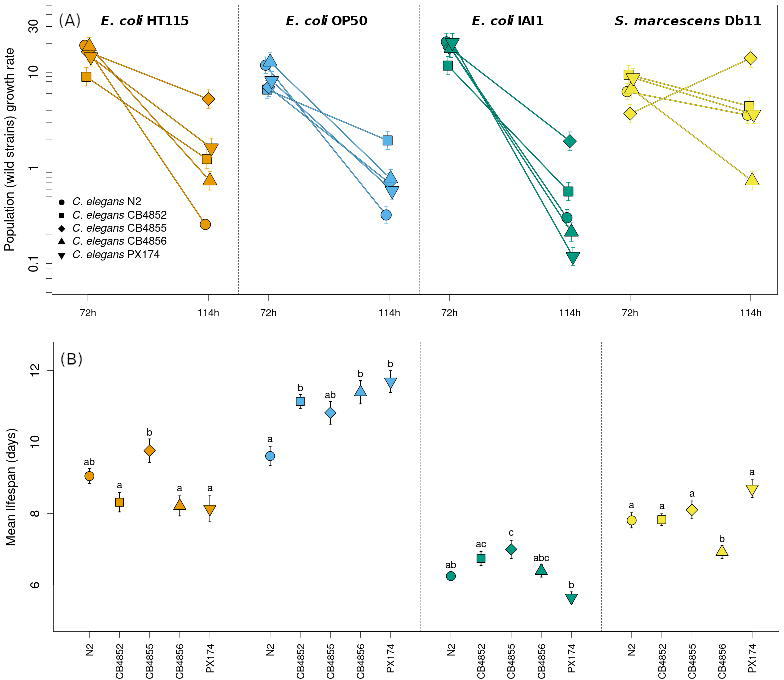
Genotype-by-environment (bacteria) interactions affect *C. elegans* population growth and survival. In (**A)**, population growth rates of the five *C. elegans* genotypes, measured at the early (72 h) and delayed reproduction (114 h) times, reveal bacterial-specific effects on the temporal dynamics of reproductive output (significant three-way interaction, *p*-value < 0.001). In (**B**), it is shown that mean lifespan also depends on the interaction between *C. elegans* genotype and bacterial strain. Letters above symbols indicate group assignment from significant post-hoc tests (*p*-value < 0.05) obtained with data for each bacteria independently. Mean estimates and SE are shown in (**A**) and predicted values are shown in (**B**). Note the logarithmic scale of the *y* axis in (**A**).

As with the population growth rates, the different bacteria also affected adult survival (Fig. 1B and Supplementary Fig. 1, χ = 629.6, df = 3, *p* <0.001) in a way that differs between the *C. elegans* strains, as revealed by a significant bacteria-host genotype interaction on lifespan (χ = 72.9, df = 12, *p* < 0.0001). Notably, one of these strains shows a departure from the expected pathogenic effects of *S. marcescens* Db11 on survival; for the PX174 genotype, lifespan in presence of S. *marcescens* (8.7 ± 0.2 days) was clearly not reduced in comparison with the one obtained with *E. coli* HT115(DE3) (8.1 ± 0.4 days). Association between population growth and lifespan was generally absent (Suppl. Table 1), apart from a marginally significant positive correlation (p-value = 0.05), obtained with *E. coli* IAI1 for population growth at 114h. Interestingly, *C. elegans* survival is markedly affected by *E. coli* IAI1 at that time (day 5), which does not happen with the other bacteria (survival of 63% in comparison with 89% to 95%). The corresponding coefficient of variation, obtained across the different genotypes, is also higher with *E. coli* IAI1 (28% in comparison with 2% to 11%).

### REPRODUCTION AND DEVELOPMENT OF A GENETICALLY DIVERSE POPULATION

Analysis of the effects of the four bacterial strains on the D00 population shows that, despite prevalent outcrossing and genetic variability, growth rate dynamics and survival are comparable to the ones observed in genetically homogeneous populations composed by single genotypes (Fig. 2). Once again, population growth rates at the early and delayed reproduction times (72h and 114h, respectively) were dependent on the bacterial strains (Fig. 2A), with a significant time-by-bacteria interaction (LRT = 2.58, df = 3, *p* < 0.001). The main effects of time (LRT = 0.25, df = 1, *p* = 0.03) and bacterial strain (LRT= 1.89, df = 3, *p* < 0.0001) were also significant. The presence of different bacteria also had significant effects on *C. elegans* survival (*p*-value < 0.0001), with lower mean lifespan observed in the presence of *E. coli* IAI1 and *S. marcescens* Db11, as expected (see Fig. 2B, adjusted *p*-values < 0.05 from pairwise comparisons are used, and Supplementary Fig. 2).

**Figure 2.**
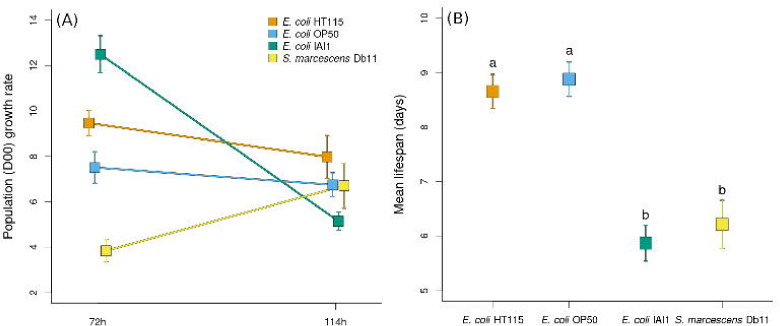
Bacterial-specific effects on the reproductive output and survival are maintained in the genetically diverse, male-female, *C. elegans* host population. As in Figure 1, (**A**) shows population growth rates measured at the early (72 h) and delayed reproduction (114 h) times, with each of the four bacteria used in this study. In (**B**), mean lifespan reveals the detrimental effects of the pathogenic *E. coli* IAI1 and *S. marcescens* Db11 bacteria in contrast with the benign *E. coli* HT115(DE3) and *E. coli* OP50 strains. Letters above symbols indicate group assignment from significant post-hoc tests (*p*-value < 0.05). In (**A**) and (**B**) mean estimates and SE are shown.

Interestingly, when some of the traits that contribute to population growth are further explored a more complex scenario is observed (Fig. 3). First, no consistently detrimental (i.e., pathogenic) effects were observed for fertility, even though it varied with the different bacteria (Fig. 3A-B). Significant differences among bacterial strains were found for lifetime fertility (*p* < 0.0001, Fig. 3A), with the highest brood size being observed with *E. coli* HT115(DE3) (371 ± 4), followed by *E. coli* OP50 (185 ± 2), *E. coli* IA1 (177 ± 2) and *S. marcescens* Db11, which resulted in a markedly reduced lifetime fertility (61 ± 1). These differences were also reflected in the reproductive schedule (Fig. 3B), as revealed by a significant time by bacteria interaction (LRT = 42.5, df = 12, *p* < 0.001). Although fertility was always maximized at day 4, the relative contribution of offspring produced before and after this peak day was dependent on the bacterial strains. For instance, with *E. coli* HT115(DE3) the higher mean estimates of fertility observed throughout the entire reproductive lifespan of the host only became significantly different from the other *E. coli* strains after day 5. In contrast, the initially diminished fertility of *S. marcescens* Db11 was no longer different from most values observed with the three *E. coli* strains from day 4 onwards (Fig. 3B). Interestingly, comparing the start of offspring production of *S. marcescens* Db11 with the ones from all *E. coli* (Fig. 3C) reveals a delay in the overall reproductive period, which could result from a specific reduction in reproductive output in the early stages or an increase in the developmental time. The comparison of developmental rates of D00 individuals with the different bacteria (Fig. 3D), indicates that the specificity of the reproductive schedule obtained with *S. marcescens* Db11 cannot be, at least fully, attributed to developmental differences. In fact, the developmental status of most *C. elegans* females in the presence of *S. marcescens* DB11 is not different from the ones observed with *E. coli* HT115 and *E. coli* OP50 (Fig. 3D). The comparison of the developmental rates obtained with the different bacteria also reveals that bacteria can have sex-specific effects (LRT= 9.63, df = 3, *p* = 0.02). Particularly, the developmental rates of males and females with *S. marcescens* DB11 are not significantly different, in contrast to what is observed with the three *E. coli* strains.

**Figure 3.**
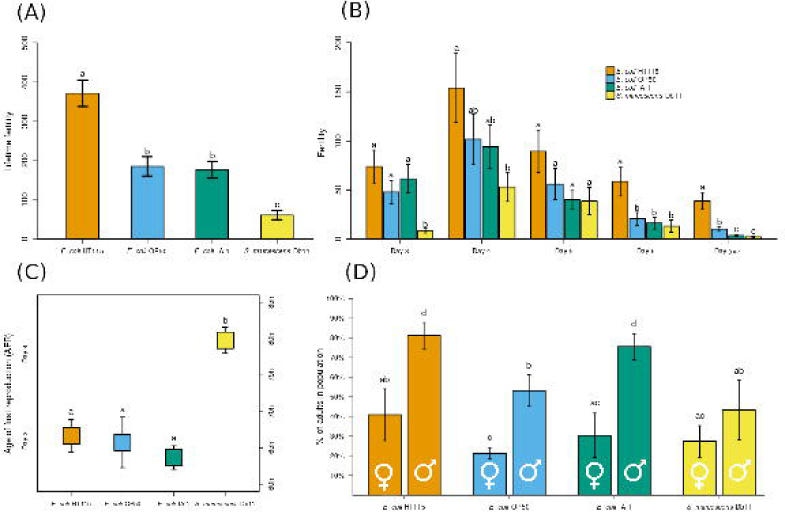
Bacteria affect the reproductive schedule and developmental rate of the genetically diverse, male-female, *C. elegans* D00 host population. (**A**) shows the lifetime reproductive success in the presence of the four different bacteria, while (**B**) shows the reproductive schedule. The age at first reproduction (AFR), given in hours and days after L1 seed (for comparison with other panels in the figure) is displayed in (**C**). In (**D**), the percentage of adult females and males at day 3 is given for the D00 population with the same bacteria. Note that, for the results shown in panels (**A**), (**B**) and (**C**), females were crossed with males that were one day older (see Methods). Means and SE are provided. Letters above bars indicate group assignment based on post-hoc tests (adjusted *p*-value < 0.05, see Methods), which in (**B**) were performed within each time period.

## Discussion

The effects of microbes on host life history raise questions about their potential role in the evolution of aging. Here, we have investigated the effects of non-pathogenic and pathogenic *E. coli* strains and of a pathogenic *Serratia marcescens* strain on the reproductive schedule and survival of *C. elegans*. Our results show that the effects of these microbes on host reproductive timing and lifespan depend on host genotype, suggesting that these traits might be subject to local adaptation to specific microbial environments in nature. We also examined a genetically diverse *C. elegans* population to study how microbial effects might affect the evolution of life-history traits, such as fertility, developmental rate, and lifespan.

### Bacteria-host genotype interactions modulate *C. elegans* reproductive timing and lifespan

We found that population growth rates and survival of five *C. elegans* genotypes varied with the presence of the three *E. coli* strains and *S. marcescens* Db11, confirming previously observed effects of bacteria on nematodes (Ikeda et al. 2007; Brooks et al., 2009; Coolon et al. 2009; Baeriswyl et al. 2010; Reinke et al. 2010; Gusarov et al. 2013; MacNeil et al. 2013; Pang and Curran 2014; Diaz et al. 2015; Dirksen et al. 2016; Samuel et al. 2016; Schulenburg and Felix 2017; Stuhr and Curran 2020; Zhang et al. 2021). We also found that these effects were host-genotype-dependent: each bacterium had a different impact on the reproductive patterns of the worms over time, which varied between genotypes, supporting the importance of host genotypes in modulating microbial effects (see Schulenburg and Ewbank 2004; Martin et al. 2017; White et al. 2019; Ekroth et al. 2021; Zhang et al. 2021).

In general, patterns of population growth revealed the typically expected decrease in offspring production after the first days of the reproductive period (Harvey and Viney 2007; Baerwysil et al. 2010; Anderson et al. 2011; Pang and Curran 2014; Carvalho et al. 2014). However, the results obtained with *S. marcescens* Db11 differed from those observed with *E. coli*. In the case of *S. marcescens* Db11, a slower decrease with age was observed, possibly indicating a reduced impact of reproductive aging or changes in self-sperm depletion. In *C. elegans,* self-sperm depletion occurs naturally due to sequential hermaphroditism in which individuals first make sperm and later switch to the production of oocytes in excess, limiting offspring numbers to the number of sperm cells produced early on (Ward and Carrel 1979; Nayak et al. 2005; Sharf et al. 2021). The differences observed in the effects of *S. marcescens* Db11 and *E. coli* strains are unlikely to be attributed solely to their pathogenicity or the stress responses they induce. If this were the case, we would expect to see similar patterns with *E. coli* IAI1 and *S. marcescens* Db11. Similarly, it is improbable that the varying amounts of bacteria present on the plates or inside the worm’s gut play a significant role in these experiments. If that were the case, the effects of the two pathogenic bacteria could differ in intensity, but not in direction. These observations suggest that other mechanisms, such as the timing of germ-line development, may be involved.

Given that bacteria can serve as both pathogens and a food source for *C. elegans* (Kim 2013; Frézal and Félix 2015; Samuel et al. 2016; Schulenburg and Félix 2017), it is possible that the differences in reproductive dynamics were due to nutritional effects. Indeed, dietary effects have been shown to change reproduction and other life-history traits of *C. elegans* (Brooks et al. 2009; Baeriwisyl et al. 2010; Reinke et al. 2010; MacNeil et al. 2013; Pang and Curran 2014; Diaz et al. 2015), e.g., interfering with the developmental timing of the worm (Baeriswyl et al. 2010; MacNeil et al. 2013; Diaz et al 2015; Stuhr and Curran 2020). Such effects, which can depend on host genotype (see Zhang et al. 2021), might explain differences in reproductive dynamics and support the importance of nutrition in shaping life-history evolution (Swanson et al, 2016).

The interaction between host genotypes and bacteria on the reproductive timing suggests that, if trade-offs between early and late reproduction are common in *C. elegans*, their evolution will likely be an important element of adaptation to microbial environments in nature. But would that translate into meaningful changes in other aging-related phenotypes, such as lifespan? In the present study, there was no evidence of trade-offs between early and late reproductive success, as there was no overall correlation observed between population growth rates and survival of *C. elegans* genotypes at different time points. Interestingly, a positive correlation was observed only in the presence of *E. coli* IAI1, where higher population growth rates at 114h were linked to increased survival. This positive association may have resulted from the lethality caused by the bacteria, leading to a demographic effect where only surviving individuals could produce offspring. This highlights the importance of condition-dependent mortality in the evolution of increased lifespan (Maklakov et al. 2015; Chen and Maklakov, 2017).

### Bacteria modulate reproduction and sex-specific development of the genetically diverse D00 population

The expectation of trade-offs in *C. elegans* reproduction has been documented in different studies involving several causes, such as self-sperm limitation in hermaphrodites, mutations affecting germline maintenance and development (Antebi 2007; Angelo and Van Gilst 2009; Maklakov and Immler 2016), or mutations in the insulin/IGF-1 signaling pathway (Gems et al. 1998; Jenkins et al. 2004; Maklakov et al. 2017). Nevertheless, some experiments focused on self-fertilizing hermaphrodites have failed to detect negative correlations between early and late-fitness related traits (Estes et al. 2005; Wu et al. 2012).

Our work with the male-female D00 population confirms that bacteria can have significant effects on trade-offs in natural systems. We observed specific bacterial effects on survival, lifetime fertility, and reproductive schedule in the presence of males and absence of selfing. Notably, the impact of bacteria on reproductive span in this population, where sperm depletion did not occur, suggests a role for bacterial effects in reproductive aging. Our results also indicate that bacterial effects on life-history traits can be sex-specific, as observed for developmental rate, and have implications for population adaptation to local microbial communities in nature. This is particularly relevant for gonochoristic (male-female) *Caenorhabditis* species like *C. remanei*, but also for *C. elegans*, despite its low expected outcrossing rates in nature (Barrière and Félix 2005; Richaud et al. 2018). Although *C. elegans* males are rarely found in nature and considered evolutionary relics with little contribution to natural populations, under challenging conditions, this may change transiently, as male frequencies and outcrossing increase during adaptation (Morran et al. 2009; Teotónio et al. 2012; Chelo and Teotónio 2013; Cutter et al. 2019).

The presence of males in *C. elegans* populations has also been found to result in a trade-off between reproduction and longevity (Wu et al. 2012; Carvalho et al. 2014). This can be attributed to sexual conflict between males and hermaphrodites, where the increased reproductive output facilitated by male sperm is countered by a reduction in the lifespan of hermaphrodites due to mating (Gems and Riddle 1996). Notably, this effect seems to occur only when self-sperm is absent, which is the case during the post-reproductive period in hermaphrodites or in mutation-derived females that lack self-sperm (Wu et al. 2012; Carvalho et al. 2014; Booth et al. 2019), as in the case of this study.

Overall, our findings underscore the major impact microbes can have on host life history. Our results suggest that selection for reproductive investment at specific times might explain microbial specificity in local adaptation, as has been observed in *D. melanogaster* (Rudman et al. 2019; Walters et al. 2020) and as has been proposed for *C. elegans* (Schulenburg and Ewbank 2004; Dirksen et al. 2016; Samuel et al., 2016; Zhang et al. 2021). This potential mechanism for adaptation is particularly relevant given the diverse, complex microbial environments that these organisms are exposed to in nature.

## Author contributions

Conception and design: Margarida Matos; Thomas Flatt and Ivo M. Chelo

Data acquisition: Josiane Santos and Ivo M. Chelo

Data analysis: Josiane Santos and Ivo M. Chelo

Data interpretation: Josiane Santos; Margarida Matos; Thomas Flatt and Ivo M. Chelo

Manuscript writing: Josiane Santos; Margarida Matos; Thomas Flatt and Ivo M. Chelo

## Acknowledgements

We are grateful to Henrique Teotónio, Sara Magalhães, and Patrícia Beldade for helpful comments on a previous version of the manuscript. Nematode strains were provided by the *Caenorhabditis* Genetics Center (CGC), funded by the NIH Office of Research Infrastructure Programs (P40 OD010440); the *E. coli* IAI1 strain by Ivan Matic; and the D00 population by Henrique Teotónio. Our research was supported by the FCT (Fundação para a Ciência e Tecnologia; grants IF/00031/2013 and PTDC/BIA-EVL/28757/2017) as well as FEDER / PORLisboa, grant LISBOA-01-0145-FEDER-028757 to I.M.C., grant SFRH/BPD/123405/2016 to J.S., and by cE3c unit funding (UIDB/00329/2021). We also acknowledge the Instituto Gulbenkian de Ciência (IGC), where the initial experiments were performed, in particular the support through the ONEIDA project (LISBOA-01-0145-FEDER-016417).

