## Supplementary Table 1 for "Microbes are potential key players in the evolution of life histories and aging in *Caenorhabditis elegans*"

**Supplementary Table 1.** Phenotypic correlations between population growth rate and lifespan of the individual genotypes used in this work.

| | *E. coli* HT115(DE3) | *E. coli* OP50 | *E. coli* IAI1 | *S. marcescens* DB11 |
|---|---|---|---|---|
| Growth 72 h – Growth 114 h | -0.28 | -0.6 | -0.34 | -0.47 |
| Growth 72 h – Lifespan | 0.41 | -0.18 | -0.4 | 0.08 |
| Growth 114 h - Lifespan | 0.14 | 0.42 | 0.88* | 0.53 |

Values are Pearson's correlation coefficients. * p-value <0.05.
