## Supplementary figures and images for "Microbes are potential key players in the evolution of life histories and aging in *Caenorhabditis elegans*"

### Supplementary Figure 1

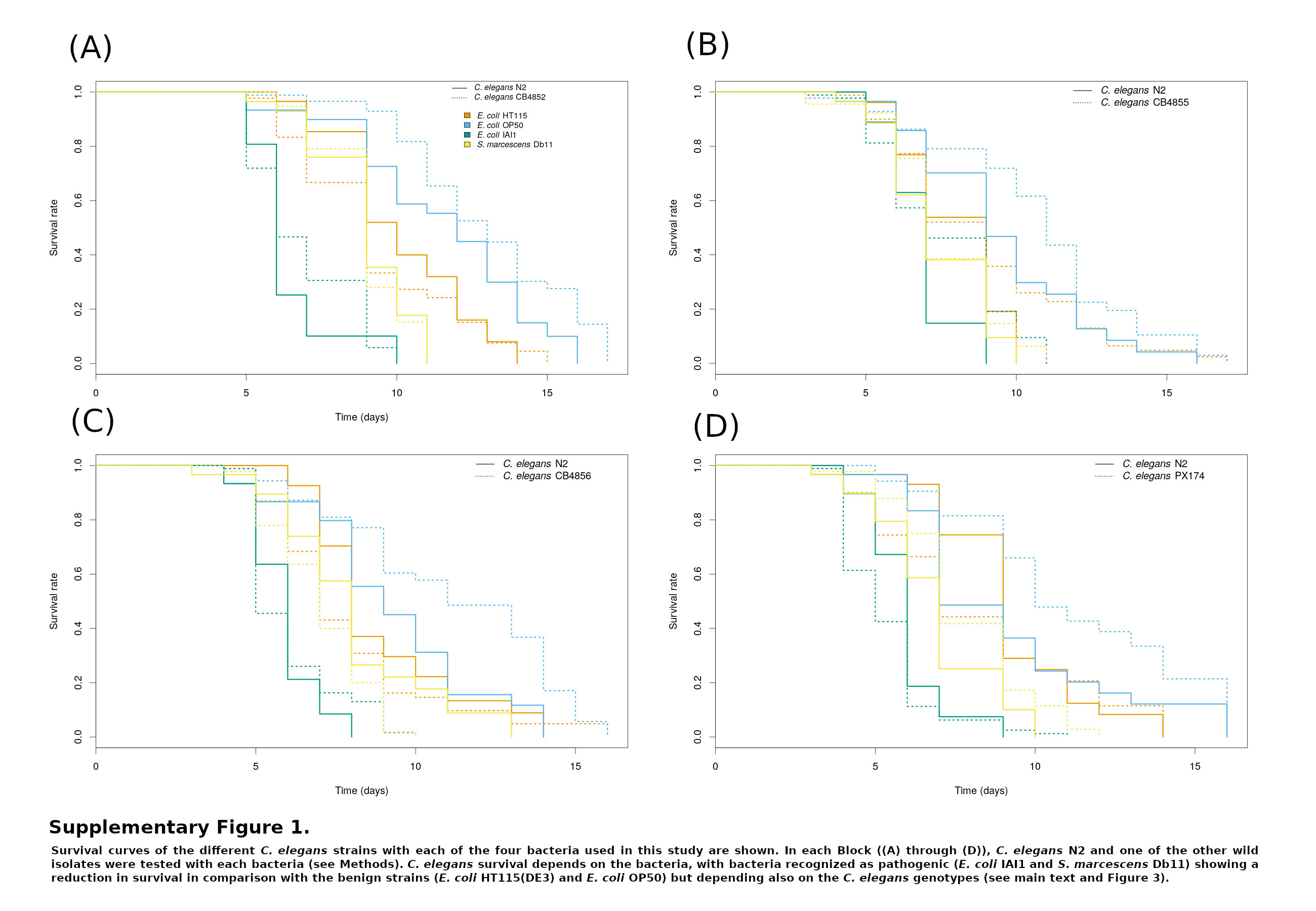

### Supplementary Figure 2

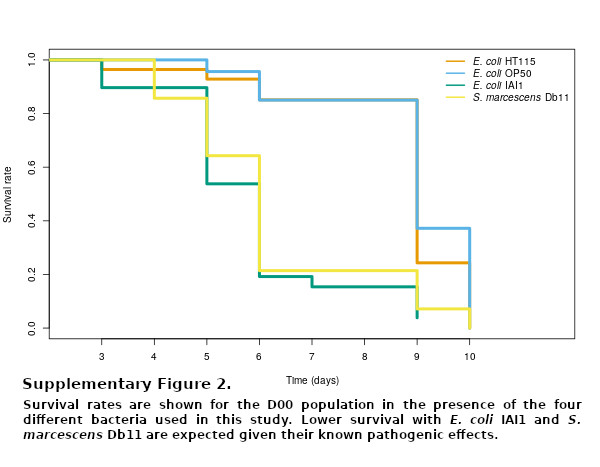
